# DUBS: A Framework for Developing <u>D</u>irectory of <u>U</u>seful <u>B</u>enchmarking <u>S</u>ets for Virtual Screening

**DOI:** 10.1101/2020.01.31.929679

**Authors:** Jonathan Fine, Matthew Muhoberac, Guillaume Fraux, Gaurav Chopra

## Abstract

Benchmarking is a crucial step in evaluating virtual screening methods for drug discovery. One major issue that arises among benchmarking datasets is a lack of a standardized format for representing the protein and ligand structures used to benchmark the virtual screening method. To address this, we introduce the Directory of Useful Benchmarking Sets (DUBS) framework, as a simple and flexible tool to rapidly created benchmarking sets using the protein databank. DUBS uses a simple input text based format along with the Lemon data mining framework to efficiently access and organize data to protein databank and output commonly used inputs for virtual screening software. The simple input format used by DUBS allows users to define their own benchmarking datasets and access the corresponding information directly from the software package. Currently, it only takes DUBS less than 2 minutes to create a benchmark using this format. Since DUBS uses a simple python script, users can easily modify to create more complex benchmarks. We hope that DUBS will be a useful community resource to provide a standardized representation for benchmarking datasets in virtual screening.

## Introduction

Small molecule protein docking is one of many essential tools applied in virtual screening pipelines for drug discovery.^1–3^ The docking protocol produces a 3D small molecule ligand pose inside the binding cavity of a protein, which is representative of the ligand conformation that is co-crystalized with the protein target. Energetics of binding site interactions of this pose are calculated to determine if the small molecule binds to the protein target, as well as, to estimate the binding affinity between the small molecule and the target. These calculations are used to virtually screen large libraries of synthetically feasible molecules to identify potential hits for specific targets.^4^

The virtual screening community has developed several benchmarking datasets to evaluate how well different docking methods perform at these vital tasks.^5–13^ One of the popular benchmarking set is the Astex Diverse Benchmark^5^ that is used to evaluate how well a given methodology can reproduce the crystal pose of a small molecule ligand in the bound (*holo*) form of a protein. Similarly, the PDBBind^14^ and related CASF^11,15^ datasets assess the ability of a docking method to produce and select a crystal-like pose for the small molecule ligand, and how well the methodology can rank the binding affinity of the small molecule. The most common metric used to identify a crystal-like pose is a root mean square deviation (RMSD) within 2.0Å of the true crystal pose^16,17^. The Directory of Useful Decoys (DUD)^8^ and Directory of Useful Decoys Enhanced (DUD-E)^9^ datasets provide decoy ligands which may not bind to the associated target proteins to evaluate docking methods that can distinguish binders from non-binders. Furthermore, the Pinc is Not Cognate (PINC)^12^ benchmark is designed to evaluate how well a methodology can ‘cross dock’ a given ligand. Cross docking refers to a process where a ligand is docked using the *holo* structure of a target protein crystalized with a different ligand. Similarly, the Holo-Apo Set 2 (HAP2)^13^ benchmark is designed to measure the performance of methods to reproduce the crystal structure of a bound ligand using the *apo* (or unbound) form of the target protein. In addition to benchmarking sets derived from known protein structures, the docking community has produced ‘challenge sets’ designed to rigorously validate a docking methodology in a blind manner. Two of these datasets are the Community Structure Activity Resource (CSAR)^18–21^ and the Drug Design Data Resource (D3R)^22–24^ using the data donated from the virtual screening community. Given the importance of these benchmarking sets as shown in the above works, it is clear that their continued development is important to improving and validating virtual screening pipelines.

The performance of any new virtual screening or docking methodology is tested on at least a few of the above benchmarks to gauge advancement in the field. Unfortunately, there are as many ways to format the input structure files for docking as there are benchmarking sets since the input file format specifications are not followed diligently. For example, the DUD^8^ and DUD-E^9^ benchmarks remove the element type for atoms specified for the target proteins (a required component of the PDB file format as per http://www.wwpdb.org/documentation/file-format.php). Additionally, the PINC^12^ benchmark only provides files in the MOL2 format where the protein atom names have been removed, causing difficulties when converting these files back to the PDB format. Finally, no reference inputs are provided for the Astex and HAP2 benchmarks and need to be rederived for any new method comparison. To support these benchmarking sets, the developers must support odd corner cases and other issues related to formatting which takes additional development time better spent improving their scientific methodology. The differences in structure of the target, decoys, and ligands results in errors associated with high quality evaluation of screening methods.^25^ In general, attempts to use a non-standard version of these benchmarks may lead to differences in reported performance compared to prior publications. To alleviate these issues, we have created the Directory of Useful Benchmarking Sets (DUBS) which aims to provide a framework for curated and standardized version of past and future benchmarking sets.

For DUBS, we chose to base our benchmarking framework on the highly standardized and widely accepted Macro Molecular Transmission Format (MMTF)^26^ for input as these files are already curated by the RCSB organization and provide a standard way of representing atomic coordinates, inter residue bonding, intra residue bonding, and other important crystallographic information. Using MMTF data structures, the entire protein data bank data takes less than 10 gigabytes of space^27^, allowing users to easily keep a copy of the PDB on their local hard drive. Furthermore, this is small enough to fit into random access memory on modern day workstations, enabling incredibly quick processing times as compared to other formats. Our recently published Lemon framework^28^ will be used for rapid processing by creating simple Python scripts for generating suitable inputs for docking software evaluation. These files are written using the Chemfiles^29^ input/output library which supports reading/writing a variety of formats in a highly standards compliant manner. We have chosen to use the PDB file format for protein input as this format is standard for use in the docking community, and the SDF file format as a preference for small molecule input as this format allows for the storage of formal charge instead of partial charge. We did not want to include the partial charges as such parameters can bias the performance of a virtual screening methodology,^30–32^ and therefore should be handled with care for each docking simulation. Similar to the input file types, DUBS can output ligands (and proteins) in a variety of formats using the Chemfiles^29^ input/output library. These file formats include SDF, Tinker XYZ, PDB, CML, ARC, CSSR, GRO, MOL2, CIF, MMTF, and SMI. DUBS can be used to standardize existing benchmarking datasets as well as rapidly create user defined new benchmarks for virtual screening applications.

## Methods

DUBS is designed to work using a specific input format that can incorporate variability in different types of information to develop user defined benchmarking sets. An overview of the DUBS pipeline is given in **Figure 1a** where an input file formatted in this specific manner is parsed using the provided python script that will be referred to as the parser, henceforth. Currently, the input format for DUBS is designed using seven tags that is used by the parser to identify different types of information in the input file. This information is stored in nine dictionaries, which keep track of alignment of each benchmark protein to a reference protein (optional), and pairing of ligands to their respective protein. To better describe the input format and options, an example ‘block’ of the input format used for DUBS is shown in **Listing 1** for hivproteasea2 from the PINC benchmarking set and the program options for each tag is provided in **Listing 2** for developing new benchmarking sets. The input files for several existing benchmarks, such as, PDBbind, Astex, PINC, HAP2, DUD-E, etc. are provided on https://github.com/chopralab/lemon/tree/dubs/ and as Supporting Information files.

**Figure 1.**
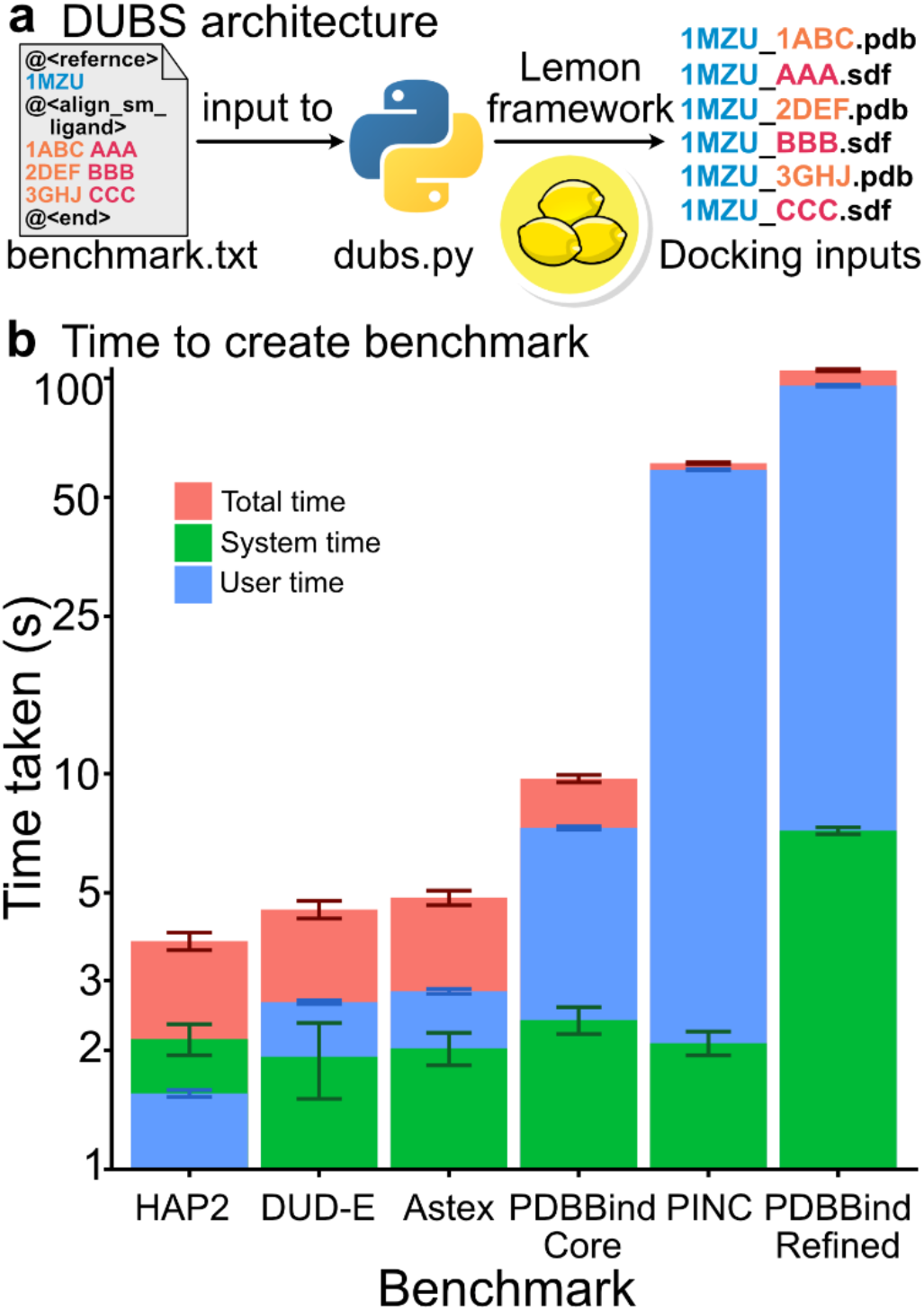
Overview of the DUBS software package is given in **(a)** where a flat text file is used to describe a given benchmarking set. A detailed description of this text file and the output of the dubs.py python script is given in the methods section. Timing results for the creation of five benchmark sets are shown in **(b)** with the number of proteins indicated.

Some benchmarking sets such as PINC and HAP2 require the use of protein alignment as they are designed to evaluate docking performance on non-native protein conformations. To address this need, a customized version of the TM-align algorithm^33^ is implemented in the Lemon framework^28^ to allow for fast and accurate alignment between crystal structures of the same target protein. Briefly, this algorithm matches residues between the reference and non-reference crystal structures and attempts to find the affine transformation which minimizes the distance between the alpha carbons of the matched residues. In contrast to the original algorithm, the Lemon implementation incorporates the chain name of the residues in addition to their standardized residue ID. Additionally, this algorithm makes use of an optimized version of the Kabsch algorithm^34^ to improve performance time.

In the input file format used by DUBS, the first tag is the reference protein tag, denoted “@<reference>”. This tag informs the parser that information on the reference protein will be read in on the next line. This line must contain the PDB ID of the reference protein and the file path of a MMTF or other Chemfiles-supported format for that protein separated by a single space. The line may also include a PDB chemical ID for the ligand bound to the reference protein so that it can be removed before alignment. This ligand code needs to be put after the file path, separate by a single space. It should be noted that only one reference protein is allowed per reference tag and the reference MMTF or other Chemfiles format file must be stored locally. The parser then saves the reference PDB ID and path as a key-value pair in the file path dictionary and the ligand to be removed in a separate dictionary with the reference PDB ID as the key. If a reference file is used, but the PDBID of the reference is not known, place a * followed by a unique number in place of the PDBID (i.e. *1, *2, *3, etc.). This will allow for proper storage of the file path and corresponding alignment information for the protein in the appropriate dictionaries.

The second tag is the aligned protein tag, denoted “@<align_prot>”. This tag informs the parser about the information on proteins that will be aligned to the reference protein and are read on multiple subsequent lines. These lines should contain the PDB IDs of any proteins planned for alignment and the PBD chemical ID of any ligands that need to be removed before alignment, separated by a single space. It should be noted that a corresponding file is not needed for these proteins. Each of these PDB IDs is stored in a list in the alignment protein dictionary with the reference PDB ID as the key. The ligands that will be removed are stored in a separate dictionary with the PDB ID of the corresponding protein as the key. As an example, this tag was used in the input file for the PINC dataset (**Listing 1**). If no PDBIDs are provided but the tag is present (**Listing 2**), DUBS will not perform alignment and the docking benchmark will be a cognate set where the ligand will be docking to its co-crystal.

The third tag and fourth tags are the small molecule (sm) and non-small molecule ligand alignment tags, denoted “@<align_sm_ligand>” and “@<align_non_sm_ligand >” respectively. These tags inform the parser that information on various proteins and their respective ligands will be read in on the subsequent lines. These lines should contain any number of paired proteins and ligands corresponding to the protein separated by a single space with only one pair per line (see **Listing 1**). For small molecule ligands, the PDB ID of the protein should be listed first on the line followed by the PDB chemical ID of the ligand, separated by a space. After parsing, each of these PDB ID’s are stored in a list in the reference dictionary with the reference PDB ID as the key. The small molecule ligand PDB chemical IDs are stored inthe small molecule ligand dictionary with their respective protein’s PDB ID as the key. For non-small molecule ligands, the subsequent lines after the tag should contain any number of paired proteins and non-small molecule ligands. The PDB ID of the protein should be listed on the first line followed by a space and then the residue code, chain ID, and residue number of the non-small molecule ligand. The PDB chemical ID code, chain ID, and residue number are separated by a dash “-” character (**Listing 2**). After parsing, each of the PDB ID’s are stored in the same list in the reference dictionary as the small molecule ligands, with the reference PDB ID as the key. The residue code, chain ID, and residue number are stored in the non-small molecule ligand dictionary with their corresponding PDB ID as the key.

The fifth is the end tag, denoted “@<end>”. This tag informs the parser that no additional information will be read and stored for the current reference protein until the next reference tag is detected. This allows for comments to be written in the file outside of each block of information at the user’s discretion. The sixth and seventh tags are no alignment small molecule and non-small molecule ligands tags, denoted by “@<no_align_sm_ligands>” and “@<no_align_non_sm_ligands>” respectively. These tags are used when no reference protein is used. In the input file, the subsequent lines should have the same format as their respective @<align_…> counterparts. If the alignment features of DUBS are to be used, the alignment tabs must appear between the reference tag and the end tag. The protein and ligand alignment tags used in between do not need to be in any order or even present at all if they are not needed. For cases where a reference protein is not used, simply put any ligands after their respective no alignment tags. If there are any user comments that need to be added in the input file, they must come before the first tag is used, or between an end tag and a reference tag. After parsing the input data and storing it in the appropriate dictionaries, the Lemon framework^28^ is used to perform alignment and scoring on the benchmarking data to produce standardized docking output files.

The DUBS software is available in Python 3.6 and gcc 6.3.0. It is freely available for installation via the Python Package Infrastructure (PyPI). All standardized formatted benchmarking sets are available for download directly from GitHub (https://github.com/chopralab/lemon/tree/dubs/). The source code for the Lemon framework API and documentation are available on GitHub at https://chopralab.github.io/lemon/latest/.

## Results and Discussion

We showcase the ability of DUBS to reproduce and standardize previously published benchmarking sets. The input file described above was created for six well established benchmarking sets that derive from the protein databank (see **Listing S2**-**S7**). We measured the amount of time required to generate these benchmarking sets when the Hadoop copy of the protein databank was stored in RAM and using a single CPU core. We generated the benchmarking inputs 50 times for each benchmarked to calculate a mean and standard deviation for each calculation. We measured the total time taken by the application as well as the ‘user’ and ‘system’ time, which represent the time spent by the application itself and the time taken accessing hardware/allocating memory, respectively. These results are shown in **Figure 1b** and indicate that DUBS is able to quickly produce a benchmarking set in less than two minutes. The most computationally expensive benchmark to generate is the PDBBind Refined set simply due to the number of ligands required to written as output files the software. The next most computationally expensive benchmark is PINC due to a large number of alignments that are required for the benchmark to be generated as there are 949 ligands as well as 60 proteins that needs to be aligned. It should be noted that the application took a greater proportion on its time in ‘user’ mode for this benchmark than in ‘system’ mode, compared to the PDBBind Refined set. These results show that DUBS can quickly recreate known benchmarking sets from the protein data bank.

In addition to the previously published benchmarks, we wanted to show how easy it is to create new benchmarking sets using the Lemon framework and DUBS. For this example benchmarking set, we created a simple script to identify all the small molecules in the PDB which interact with a Heme group (see original Lemon publication^28^ for details). This script identified 1974 complexes that matches this criterion and created a DUBS input file without reference alignment. Of these complexes, 904 of them have unique small molecule ligands and we selected the complex with the lowest crystal structure resolution as representative of the small molecule/Heme protein complex. All steps in this methodology took less than 10 minutes, showing the power of the Lemon and DUBS pipeline. Depending on the user several other complex but standardized benchmarking sets can be created using DUBS for future virtual screening applications.

## Conclusion

In the work, we have presented a new software package, DUBS, for the creation of standards compliant benchmarking sets. DUBS uses a simple, flat text file as input to create docking input files for existing and new benchmarking sets for use in virtual screening within five minutes on modern hardware. Using this file format, we have reproduced 6 popular benchmarking sets and created an example of a new benchmarking set that can be used to evaluate how well docking methods can reproduce the binding mode of a compound in the presence of a Heme group. We believe that the DUBS framework will enable users to create new customizable and standardized benchmarking sets rapidly to foster a new era of useful virtual screening pipelines.

~~~
hivproteasea2
@<reference>
1MTB pinc_reference_files/hivproteasea2/1MTB.mmtf HPH
@<align_prot>
4HVP 2NC
2FGV NTB
1MTR PI6
2AID THK
@<align_sm_ligands>
3MXD K53
2FGU NTB
3EM3 478
3EM6 017
3OY4 017
3EKV 478
@<end>
~~~

**Listing 1.** This is an example of the input format used for DUBS. This example is hivproteasea2 from the PINC benchmarking set.

**a** DUBS input format when a reference protein is used

~~~
Comments go here
@<reference>^1^
PDB_ID^3^ reference_file_path^3^ PBD_chemical_ID^4^
@<align_prot>^2^
PDB_ID^3^ PBD_chemical_ID^3^
@<align_sm_ligands>^2^
PDB_ID^3^ PDB_chemcial_ID^3^
@<align_non_sm_ligands>^2^
PDB_ID^3^ residue_code-chain_ID-residue_number^3^
@<end>^1^
Additional comments here
~~~

**b** DUBS input format when a reference protein is not used

~~~
Comments go here
@<no_align_sm_ligands>^2^
PDB_ID^3^ PDB_chemical_ID^3^
@<no_align_non_sm_ligands>^2^
PDB_ID^3^ residue_code-chain_ID-residue_number^3^
@<end>^2^
Additional comments here
~~~

**Listing 2**. Showing required and optional formatting arguments for both input files (a) involving a reference protein and (b) not involving a reference protein. Denoted are required tags (1), optional tags

(2), required tag elements (3), and optional tag elements (4).

## Acknowledgments

This work was supported in part by a Purdue University start-up package from the Department of Chemistry at Purdue University, Ralph W. and Grace M. Showalter Research Trust award, the Integrative Data Science Initiative award, the Jim and Diann Robbers Cancer Research Grant for New Investigators award and NIH NCATS ASPIRE Design Challenge awards to Gaurav Chopra, a Lynn Fellowship to Jonathan Fine and a Ross Fellowship to Matthew Muhoberac. Additional support, in part by, a NCATS Clinical and Translational Sciences Award from the Indiana Clinical and Translational Sciences Institute (UL1TR002529), and the Purdue University Center for Cancer Research NIH grant P30 CA023168 are also acknowledged. The content is solely the responsibility of the authors and does not necessarily represent the official views of the National Institutes of Health.

## TOC Figure

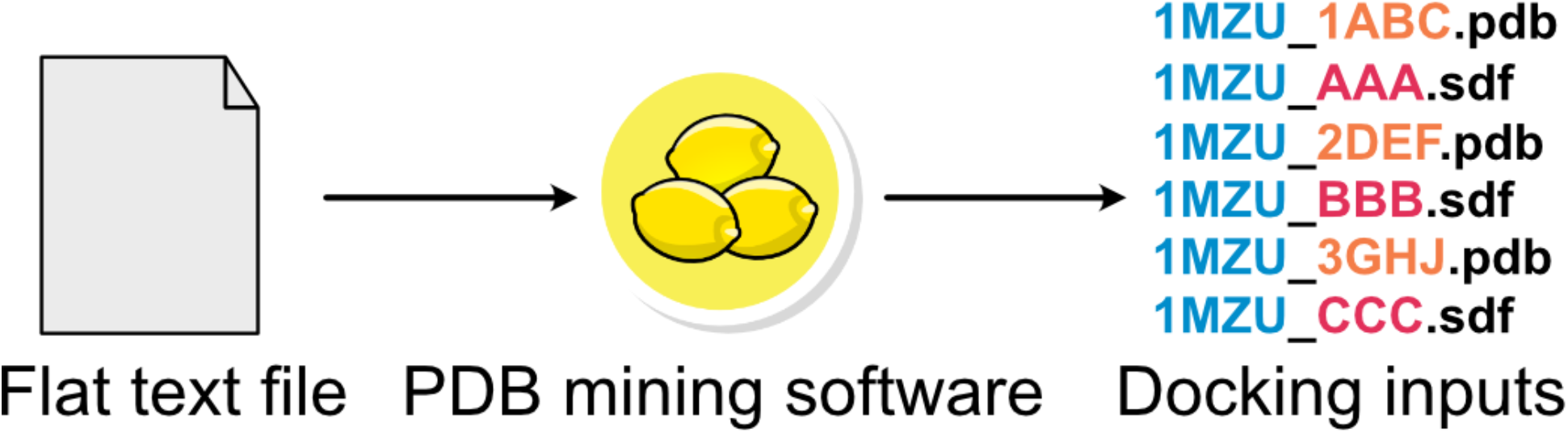

