## Supporting Information for "DUBS: A Framework for Developing <u>D</u>irectory of <u>U</u>seful <u>B</u>enchmarking <u>S</u>ets for Virtual Screening"

DUBS: A Framework for Developing Directory of Useful Benchmarking Sets for Virtual Screening

### **Supporting Information**

**Listing 1: DUBS python script**

```
from candiy_lemon import lemon
import sys

# List of dictionaries to keep track of parts of the file

# Key: reference pdbid, Value: path to .mmtf file
pathDict = {}
# Key: reference pdbID, Value: chemical ID of ligand bound to reference protein for removal
referenceLigandDict = {}
# Key: reference pdbID, Value: list of sm or non-sm protein
referenceDict = {}
# Key: reference pdbID, Value: list of proteins to aling to reference (like in pinc)
alignProtDict = {}
# Key: align protein pdbID, Value: ligand chemical ID code for ligand removal
alignProtLigandDict = {}
# Key: pdbID, Value: chemical id for SM ligand
pdbIDSMDict = {}
# Key: pdbID, Value: tuple(resCode, chainID, residue ID)
pdbIDNonSMDict = {}
# Key: pdbID, Value: chemical id for SM ligand
noAlignSMDict = {}
# Key: pdbID, Value: tuple(resCode, chainID, residue ID)
noAlignNonSMDict = {}

entries = lemon.Entries()

# Method for parsing a formated input file
def parse_input_file(fname):
    # Open file and initialize flags to 0
    f = open(fname,"r")
    curRefPdbID = ""
    refFlag = 0
    protFlag = 0
    SMLigFlag = 0
    nonSMligFlag = 0
    noAlignSMFlag = 0
    noAlignNonSMFlag = 0

    for line in f:
        # Check to see if the line contains any of the tags
        # Set appropriate flags if it does
        if line.startswith("@<reference>"):
            refFlag = 1
            protFlag = 0
            SMLigFlag = 0
            nonSMligFlag = 0
            noAlignSMFlag = 0
            noAlignNonSMFlag = 0
        elif line.startswith("@<align_prot>"):
            refFlag = 0
            protFlag = 1
            SMLigFlag = 0
            nonSMligFlag = 0
            noAlignSMFlag = 0
            noAlignNonSMFlag = 0
        elif line.startswith("@<align_sm_ligands>"):
            refFlag = 0
            protFlag = 0
            SMLigFlag = 1
            nonSMligFlag = 0
            noAlignSMFlag = 0
            noAlignNonSMFlag = 0
        elif line.startswith("@<align_non_sm_ligands>"):
            refFlag = 0
            protFlag = 0
            SMLigFlag = 0
            nonSMligFlag = 1
            noAlignSMFlag = 0
            noAlignNonSMFlag = 0
        elif line.startswith("@<no_align_sm_ligands>"):
            refFlag = 0
            protFlag = 0
            SMLigFlag = 0
            nonSMligFlag = 0
            noAlignSMFlag = 1
            noAlignNonSMFlag = 0
        elif line.startswith("@<no_align_non_sm_ligands>"):
            refFlag = 0
            protFlag = 0
            SMLigFlag = 0
            nonSMligFlag = 0
            noAlignSMFlag = 0
            noAlignNonSMFlag = 1
        elif line.startswith("@<end>"):
            refFlag = 0
            protFlag = 0
            SMLigFlag = 0
            nonSMligFlag = 0
            noAlignSMFlag = 0
            noAlignNonSMFlag = 0
        else:
            # If the line does not contain a flag
            # Add info to appropriate dictionary based of set flags
            if refFlag == 1:
                pdbID = line.split(" ")[0].strip().upper()
                path = line.split(" ")[1].strip()
                curRefPdbID = pdbID
                pathDict[pdbID] = path

                if len(line.split(" ")) == 3:
                    chemID = line.split(" ")[2].strip().upper()
                    referenceLigandDict[pdbID] = chemID

            elif protFlag == 1:
                pdbID = line.split(" ")[0].strip()
                chemID = line.split(" ")[1].strip()

                if alignProtDict.get(curRefPdbID,0) == 0:
                    alignProtDict[curRefPdbID] = [pdbID]
                else:
                    alignProtDict[curRefPdbID].append(pdbID)

                if alignProtLigandDict.get(pdbID, 0) == 0:
                    alignProtLigandDict[pdbID] = [chemID]
                else:
                    alignProtLigandDict[pdbID].append(chemID)

                entries.add(pdbID)

            elif SMLigFlag == 1:
                pdbID = line.split(" ")[0].strip()
                chemID = line.split(" ")[1].strip()

                if referenceDict.get(curRefPdbID,0) == 0:
                    referenceDict[curRefPdbID] = [pdbID]
                else:
                    referenceDict[curRefPdbID].append(pdbID)

                if pdbIDSMDict.get(pdbID,0) == 0:
                    pdbIDSMDict[pdbID] = [chemID]
                else:
                    pdbIDSMDict[pdbID].append(chemID)

                entries.add(pdbID)

            elif nonSMligFlag == 1:
                pdbID = line.split(" ")[0].strip().upper()
                residueCode = line.split(" ")[1].split("-")[0].strip().upper()
                chainID = line.split(" ")[1].split("-")[1].strip().upper()
                residueID = line.split(" ")[1].split("-")[2].strip().upper()

                if referenceDict.get(curRefPdbID,0) == 0:
                    referenceDict[curRefPdbID] = [pdbID]
                else:
                    referenceDict[curRefPdbID].append(pdbID)

                if pdbIDNonSMDict.get(pdbID,0) == 0:
                    pdbIDNonSMDict[pdbID] = [tuple([residueCode,chainID,residueID])]
                else:
                    pdbIDNonSMDict[pdbID].append(tuple([residueCode,chainID,residueID]))

                entries.add(pdbID)

            elif noAlignSMFlag == 1:
                pdbID = line.split(" ")[0].strip().upper()
                chemID = line.split(" ")[1].strip().upper()

                if noAlignSMDict.get(pdbID,0) == 0:
                    noAlignSMDict[pdbID] = [chemID]
                else:
                    noAlignSMDict[pdbID].append(chemID)

                entries.add(pdbID)

            elif noAlignNonSMFlag == 1:
                pdbID = line.split(" ")[0].strip().upper()
                residueCode = line.split(" ")[1].split("-")[0].strip().upper()
                chainID = line.split(" ")[1].split("-")[1].strip()
                residueID = line.split(" ")[1].split("-")[2].strip()

                if noAlignNonSMDict.get(pdbID,0) == 0:
                    noAlignNonSMDict[pdbID] = [tuple([residueCode,chainID,residueID])]
                else:
                    noAlignNonSMDict[pdbID].append(tuple([residueCode,chainID,residueID]))

                entries.add(pdbID)

# Get from the command line
# for testing we also can hard set the path
if len(sys.argv) > 4:
    input_file_path = sys.argv[1]
    hadoop_path = sys.argv[2]
    cores = int(sys.argv[3])
    outdir = sys.argv[4]
else:
    #TODO change this if needed for testing
    print("You must give the input file, path to RCSB hadoop files, number of cores, and working directory")
    sys.exit(1)

# Define Lemon workflow class
class MyWorkflow(lemon.Workflow):
    def __init__(self):
        lemon.Workflow.__init__(self)

        self.reference_structures = {}
        for key, value in pathDict.items():
            self.reference_structures[key] = lemon.open_file(value)

        self.noAlignSMDict = noAlignSMDict

        self.outdir = outdir

        self.write_all_proteins = False

    def worker(self, entry, pdbid):
        # Define and assign the reference pdbid
        refpdbid = ""
        # mode is 0 unassigned, 1 for alignment for protein, 2 for alignment for ligand
        mode = 0

        # Check for pdbID as a protein to be aligned (like in PINC)
        for key, value in alignProtDict.items():
            if pdbid in value:
                refpdbid = key
                mode = 1

        # Check for protein-ligand pair for alignment
        for key, value in referenceDict.items():
            if pdbid in value:
                refpdbid = key
                mode = 2

        if mode == 0:
            for ligand_code in self.noAlignSMDict.get(pdbid, []):
                rns = lemon.ResidueNameSet()
                rns.append(lemon.ResidueName(ligand_code))
                ligand_ids = lemon.select_specific_residues(entry, rns)
                lemon.prune_identical_residues(entry, ligand_ids)

                for ligand_id in ligand_ids:
                    protein = lemon.Frame()
                    ligand = lemon.Frame()

                    lemon.separate_protein_and_ligand(entry, ligand_id, 25.0, protein, ligand)
                    lemon.write_file(protein, self.outdir + "/" + pdbid + "_" + ligand_code + ".pdb")
                    lemon.write_file(ligand, self.outdir + "/" + pdbid + "_" + ligand_code + ".sdf")

            return pdbid + " no alignment\n"

        elif mode == 1:
            # If we need to align to a protein (like in PINC)
            alignment = lemon.TMscore(entry, self.reference_structures[refpdbid])
            positions = entry.positions()
            lemon.align(positions, alignment.affine)

            lemon.write_file(entry, self.outdir + "/" + refpdbid + "_" + pdbid + ".pdb")

            return "Align Protein: " + refpdbid + "_" + pdbid + " to " + refpdbid + " with score of " + str(alignment.score) + "\n"

        elif mode == 2:
            # If we are doing ligand alignment, that can be done here
            # Get a list of the ligands associated with the protein we are trying to align
            SM_ligandList = pdbIDSMDict.get(pdbid, [])
            Non_SM_ligandList = pdbIDNonSMDict.get(pdbid, [])

            alignment = lemon.TMscore(entry, self.reference_structures[refpdbid])
            positions = entry.positions()
            lemon.align(positions, alignment.affine)

            if len(SM_ligandList) > 0:

                for ligand_code in SM_ligandList:

                    rns = lemon.ResidueNameSet()
                    rns.append(lemon.ResidueName(ligand_code))

                    ligand_ids = lemon.select_specific_residues(entry, rns)
                    lemon.prune_identical_residues(entry, ligand_ids)

                    for ligand_id in ligand_ids:
                        protein = lemon.Frame()
                        ligand = lemon.Frame()

                        lemon.separate_protein_and_ligand(entry, ligand_id, 25.0, protein, ligand)
                        lemon.write_file(ligand, self.outdir + "/" + refpdbid + "_" + pdbid + "_" + ligand_code + ".sdf")

                        if self.write_all_proteins:
                            lemon.write_file(protein, self.outdir + "/" + refpdbid + "_" + pdbid + "_" + ligand_code + ".pdb")

            return "Align Protein: " + pdbid + " to " + refpdbid + " with score of " + str(alignment.score) + "\n"

    def finalize(self):
        pass

# Parse the input file
parse_input_file(input_file_path)
print(pathDict)
# Initilize the workflow
wf = MyWorkflow()

# TODO Get these from the command-line or ask the user
lemon.launch(wf, hadoop_path, cores, entries)
```

**Listing 2: Input for Astex dataset**

@<no_align_sm_ligands>

1m2z DEX

1pmn 984

1s3v TQD

1t40 ID5

1ke5 LS1

1p2y NCT

1hww SWA

1oyt FSN

1hp0 AD3

1tow CRZ

1xoz CIA

1mmv 3AR

1z95 198

1r1h BIR

1hq2 PH2

1meh MOA

1unl RRC

2br1 PFP

1y6b AAX

1xoq ROF

1n46 PFA

1ywr LI9

1x8x TYR

1n2v BDI

1w2g THM

1yqy 915

1g9v RQ3

1ygc 905

1k3u IAD

1sg0 STL

1gpk HUP

1hwi 115

1yvf PH7

1tz8 DES

1gm8 SOX

1u1c BAU

1oq5 CEL

1of6 DTY

1lpz CMB

1q41 IXM

1n1m A3M

1kzk JE2

1lrh NLA

1r58 AO5

1v48 HA1

2bm2 PM2

1gkc NFH

1t9b 1CS

1jla TNK

1xm6 5RM

1v0p PVB

1nav IH5

1s19 MC9

1w1p GIO

1opk P16

1l7f BCZ

1vcj IBA

1r55 097

1u4d DBQ

1owe 675

1jd0 AZM

1ig3 VIB

1tt1 KAI

1jje BYS

1l2s STC

1ia1 TQ3

2bsm BSM

1hnn SKF

1j3j CP6

1v4s MRK

1sq5 PAU

1sqn NDR

1uml FR4

1p62 GEO

1yv3 BIT

1t46 STI

1q1g MTI

1uou CMU

1mzc BNE

1r9o FLP

1q4g BFL

1n2j PAF

1hvy D16

1of1 SCT

1sj0 E4D

**Listing 3: Input for PDB Bind Core dataset**

@<no_align_sm_ligands>

2brb PFQ

2v00 V15

2qe4 JJ3

3jvs AGY

5aba UL7

1ps3 KIF

3coz 54H

3qqs 17C

2iwx M1S

4eor 4SP

2w66 HQ6

3cj4 SX5

3d6q U3S

2zcr B69

3e93 19B

4agp P51

4djv 0KM

1oyt FSN

4mme 29Q

3utu 1TS

2al5 FWD

4j28 EAT

2zda 32U

4ea2 RWZ

4qd6 30T

3wz8 IXV

1lpg IMA

2pog WST

2xdl 2DL

2xj7 GC2

3syr GPK

1r5y DQU

1z95 198

1qkt EST

3b5r B5R

3d4z GIM

2zb1 GK4

2w4x STZ

2fxs RDA

1z6e IK8

3gc5 2MQ

4ddk 0HN

3ui7 C1L

3gbb MS8

4llx 5ZE

4w9h 3JF

2wtv ZZL

3rlr 3RR

2br1 PFP

2wer RDC

3lka M4S

4dli IRG

3gnw XNC

2zcq B65

3gv9 GV9

3wtj TH4

3u9q DKA

3ary I5I

3fv2 NDZ

2cbv CGB

3pww ROC

4m0z M0Z

4kzq DFL

3b27 B2T

3mss MS7

4eo8 0S3

1u1b PAX

4de3 DN8

3kgp 4AZ

2xbv XBV

2p4y C03

3uex STE

1ydt IQB

1h22 E10

1o0h ADP

1q8t Y27

3ozs OZS

3pyy 3YY

4qac KK3

3oe4 610

2xys SY9

3bgz VX3

3coy 53H

4e5w 0NT

4abg 91B

2v7a 627

4jia 1K3

3fcq M3S

2xii TA9

4cra XJ8

1bzc TPI

4twp AXI

1nc1 MTH

3bv9 DAR

2qnq QN3

4e6q 0NV

1owh 239

4ciw XH2

2qbr 910

1mq6 XLD

4k18 1OB

4lzs L46

5dwr 5H7

1c5z BAM

2zy1 830

1o5b ESI

4f2w TDI

2qbq 4B3

3ag9 A02

4f09 JAK

4agq P96

3gr2 GF4

3u8k 09P

3f3d MET

4w9c 3JG

1gpk HUP

3myg EML

3ejr HN4

4pcs 2M7

1gpn HUB

2hb1 512

4crc OTM

2r9w 23C

2ymd SRO

2yfe YFE

3l7b DKZ

3dx2 MZB

4hge 15V

1vso AT1

3ivg FG5

3g2n OAK

4dld TZG

1qf1 TI1

1ydr IQP

3n7a FA1

3uo4 0C0

3cyx ROC

4gid 0GH

3f3c PFF

3zt2 ZT2

2wnc TKT

1p1n KAI

3ehy TBL

3g2z GZ2

3aru PNX

3ge7 AFQ

3g0w LGB

4agn NXG

2y5h Y5H

3ueu DAO

2x00 GYN

4cig X0P

3udh 091

1nvq UCN

2wvt FHN

3gy4 PBZ

3nw9 637

3qgy PQC

4w9i 3JS

1p1q AMQ

4ddh MS0

3nq9 OCA

3o9i A61

3prs RIT

3fur Z12

1h23 E12

4ivc 1J6

3rr4 HRD

5c28 4XV

3ryj RYJ

4gr0 R4B

4ivd 15T

4kz6 ZB6

2j7h AZF

4rfm 3P6

3ozt OZZ

2fvd LIA

3uuo 0CV

3kr8 XAV

4f3c BIG

3pxf 2AN

3ao4 833

4eky D1J

1z9g RRT

2vkm BSD

3zdg XRX

3uew PLM

4jxs 18U

3jya LWG

3arv SAU

2wbg LGS

1sqa UI1

4gfm 0X2

4de2 DN3

4m0y M0Y

2xnb Y8L

3f3e LEU

3nx7 NHK

3n76 CA2

3e5a VX6

3u8n 09S

2vvn NHT

3fv1 DYH

4ty7 39F

1o3f 696

4f9w LM4

5c2h 4XU

3f3a TRP

2c3i IYZ

2p15 EZT

1eby BEB

1uto PEA

3up2 0C8

4bkt QD0

1syi CPW

2vw5 BC6

1q8u H52

3acw 651

3b1m KRC

4w9l 3JJ

3dxg U5P

3uev MYR

1k1i FD1

3oe5 611

5a7b KMN

3g31 GF1

1s38 MAQ

1bcu PRL

2cet PGI

3ebp CPB

4u4s 3C1

3b68 B68

4k77 1Q4

1y6r MTM

3r88 14F

3uri PRO

3p5o EAM

4j3l AJ5

1nc3 FMC

3arq DM5

3rsx RSV

3arp DEQ

3zso O2N

2j78 GOX

4j21 AJ6

3tsk QEG

3zsx N44

2yge GMY

3jvr AGX

1w4o UA3

2xb8 XNW

3e92 G6A

2qbp 527

4x6p 3YU

1e66 HUX

3dd0 EZL

4de1 0J6

4wiv 3P2

4owm 3F0

4ih5 12R

1yc1 4BC

4ih7 1ER

3twp SAL

1pxn CK6

2weg FBV

4mgd 27N

4ivb 1J5

4ogj 2TA

2wn9 ZY5

3n86 RJP

2wca NP6

3k5v STJ

4gkm 683

3b65 3B6

4jfs 16Z

4kzu A73

3dx1 YHO

3kwa SPM

2yki YKI

3u5j 08H

4jsz FB2

4cr9 OTW

**Listing 4: Input for PINC dataset**

hivproteasea2

@<reference>

1MTB pinc_reference_files/hivproteasea2/1MTB.mmtf HPH

@<align_prot>

4HVP 2NC

2FGV NTB

1MTR PI6

2AID THK

@<align_sm_ligands>

3MXD K53

2FGU NTB

3EM3 478

3EM6 017

3OY4 017

3EM4 DR7

3EKT 017

2QHY MZ1

3GI5 K62

2QI3 MZ5

3O9H K2E

3EL0 1UN

3I7E DJR

2QI1 MZ4

3EKP 478

3EL4 ROC

3EKX 1UN

2QI0 MZ3

2QI4 MZ6

3O9F K2D

3EKW DR7

3O9I A61

2QHZ MZ2

3EL1 DR7

3MXE K54

3EKY DR7

3OXW 017

2Q3K MUW

3GI4 K60

2QI6 MZ8

3O9G F53

3EKQ ROC

3EL9 DR7

3OXX DR7

3GI6 D78

3EL5 1UN

2QI7 MZ9

2QI5 MZ7

3R4B 74T

3EKV 478

@<end>

hivproteasebr

@<reference>

1A8K pinc_reference_files/hivproteasebr/1A8K.mmtf 0Q4

@<align_prot>

1IIQ 0ZR

2UPJ U02

1D4Y TPV

1AAQ PSI

@<align_sm_ligands>

1MSN JE2

1MRX K57

2QNN QN1

2O4P TPV

2O4L TPV

1MRW K57

2PQZ G0G

2O4S AB1

2QD7 065

2QNQ QN3

1Z8C 0ZS

2PWR G4G

2QNP QN2

1RL8 RIT

2HC0 AB2

3QPJ DTD

1ZLF 0ZR

1XL5 190

3QRM DTD

3QRO DTD

2PWC G3G

3BHE BZN

2PK5 075

1XL2 189

1IZI Q50

3UCB 017

3KDB 006

3QBF JHG

1ZPK 0ZS

3NLS 016

3QPJ N4I

2O4K DR7

1ZBG 0ZS

2BB9 AKC

3QOZ 017

2R43 G3G

3QRS NK8

1M0B 0ZQ

2QHC AB1

3JVY 017

3CKT YDP

3QRM NK7

3QRO NK9

2P3B 3TL

3UFN ROC

3JVW DMP

@<end>

hsp90alpha

@<reference>

2H55 pinc_reference_files/hsp90alpha/2H55.mmtf DZ8

@<align_prot>

1UYG PU2

1YC4 43P

2UWD 2GG

1YET GDM

@<align_sm_ligands>

3HYZ 42C

2YEJ ZZ3

3FT5 MO8

4EGK RDC

2WI3 ZZ3

3QTF 05S

2JJC LGA

3BM9 BXZ

3HZ1 37D

3R91 06H

3HYY 37D

2XDL 2DL

2YKJ YKJ

2WI2 ZZ3

3EKR PY9

3R4O FU3

2YEA 2A9

3B26 B2L

2XDU LGA

2YEB 2K4

4EFU EFU

2XHR C0P

3OW6 MEX

3HZ5 Z64

3B25 B2K

3FT8 MOJ

2XHX T5M

3RKZ 06T

3VHD VHE

3INW JZB

4EGI B2J

3EKO PYU

2WI5 ZZ5

2YEC XQ0

2VCJ 2EQ

3HZ1 42C

2XDK XDK

3R4P FU7

2QG2 A91

2YI6 6QM

3K97 4CD

2XHT C0Y

3B28 B2X

3OWD MEY

2YKI YKI

3K99 PFT

2YEJ XQK

3R4N FU5

2YJW YJW

3K98 1RC

2VCI 2GJ

2YE2 XQI

2XDX WOE

2XJJ L81

2WI6 ZZ6

3R4M WOE

2YED ADE

2YE4 2FY

2YEE 2EC

2YE3 VXX

2YE6 2AE

2YE7 2GA

2YE5 2A7

3B24 B2J

2XAB VHD

2XJX XJX

3RLR 3RR

2YE8 2D3

2YEH 2KU

3HHU 819

2YEI XQI

3BMY CXZ

2XJG XJG

2YE9 2D4

4EFT EFT

2YI0 YI0

2YK2 YJW

3VHA VHA

3RLP 3RP

3RLQ 3RQ

2WI4 ZZ4

2YI7 BZ8

3D0B SNX

3QDD 94M

4EGH 0OY

3MNR SD1

4EEH HH6

3INX JZC

3VHC VHC

3O0I P54

2YEI XQK

2YKC YKC

2YKB YKB

3R92 06J

2WI7 2KL

3OWB BSM

3B27 B2T

3HEK BD0

@<end>

pparg

@<reference>

2ATH pinc_reference_files/pparg/2ATH.mmtf 3EA

@<align_prot>

2I4J DRJ

1FM6 BRL

2HWQ DRY

1FM9 570

@<align_sm_ligands>

3ET3 ET1

3CWD LNB

3FUR Z12

3ET0 ET0

3AN3 M7S

3G9E RO7

2POB GW4

2P4Y C03

3BC5 ZAA

3B3K LRG

3OSW XDI

2Q61 SF1

3U9Q DKA

3OSI XDH

2Q6S PLB

3PO9 XPT

3D6D LRG

3R5N MLO

3KMG 538

2Q5P 241

2VSR 9HO

3FEJ CTM

3R8I XCX

3AN4 M7R

3V9Y 24L

3V9T 17L

2VST 243

3VN2 TLS

3PBA ZXG

2ZK6 C08

3S9S M0T

3V9V 21L

3TY0 082

2Q6R SF2

3CWD LNA

3CDS GRR

3IA6 UNT

3LMP CEK

2XKW P1B

3ADT HID

3CS8 BRL

2Q59 240

2OM9 AJA

3T03 3T0

3R8A HIG

3B1M KRC

2YFE YFE

3QT0 486

3K8S Z27

2Q5S NZA

3ADS IMN

3ADV SRO

2Q8S L92

3ADU MYI

3CDP YRG

3H0A D30

3B0Q MC5

3ADW MYI

3GBK 2PQ

2I4Z DRH

3ADX IMN

3NOA 5BC

@<end>

ptp1b

@<reference>

1KAK pinc_reference_files/ptp1b/1KAK.mmtf FNP

@<align_prot>

1NL9 989

1C85 OBA

1G7G INX

1ECV 878

@<align_sm_ligands>

2FJM 073

2CMA F20

3CWE 825

2ZN7 410

2BGD T1D

2BGE T2D

2QBS 024

2B07 598

1WAX LO1

2CNF F32

1ONZ 968

2F6V SK2

1ONY 588

2CMB F17

2F71 UN7

2CMC DFM

2AZR 982

2CM7 IZD

2F6Y ENT

2QBQ 4B3

1Q6T 600

1Q6P 213

2H4K 509

2H4G 694

2VEU IZ1

1Q1M 234

2HB1 512

2F70 UN6

2F6W UN3

2CM8 F16

2FJN 073

1PH0 418

1Q6J 335

1Q6S 214

1PYN 941

2F6T 1C2

2CNE DFJ

1NZ7 901

1Q6N P90

2ZMM 35B

2VEY IZ5

2QBR 910

1QXK 429

2CNG IZE

2CNI IZF

2VEV IZ2

2NTA 521

1XBO IX1

2CNH IZB

2F6Z UN5

1PXH SNA

1Q6M P27

@<end>

betasecretase1

@<reference>

2IQG pinc_reference_files/betasecretase1/2IQG.mmtf F2I

@<align_prot>

2G94 ZPQ

1W51 L01

2Q11 XX4

2OHQ 7IP

@<align_sm_ligands>

3BUF AEG

3CKP 012

3TPR 5HA

3BUG AEH

3CKR 009

2WF3 ZY3

3NSH 957

3LNK 74A

3RSX RSV

4DPI 0N1

3PI5 3P5

4DH6 0KN

3DM6 757

2ZE1 411

3H0B B35

3IXJ 586

3S7L 591

4DJY 0KR

3IVI 2LI

2WJO QUD

3L58 CS5

2XFJ VG5

2VJ6 VG5

4DJU 0KK

3RTH RTH

3UDY 09G

3RU1 3RU

3N4L 842

3INH 569

4DJV 0KM

2WEZ ZYE

3UDN 09B

3I25 MV7

2VJ7 VG6

3RVI RVI

3UDP 09D

4DJW 0KP

3SKG PB8

3UFL 508

3K5C 0BI

3DUY AFJ

3KMX G00

3DV1 AR9

2XFI XFI

3LPI Z74

2VJ9 VG7

3IN3 472

3LPJ Z75

3IND 593

2WF1 ZY1

3L5F BDX

3QBH QBH

2VNN CM7

2VNM CM8

3CIB 314

3HVG EV0

3IGB 454

3INE X17

2ZJN F1N

3RTN RTN

2VIY VG3

3CID 318

3LHG Z81

2VIZ VG4

3INF X45

3DV5 BAV

3IN4 BX2

3CIC 316

2WF4 ZY4

3MSJ EV3

3IVH 1LI

3L5E BDW

3S7M 532

4ACX S8Z

3UDJ 092

3UDH 091

2WF2 ZY2

3U6A 18P

4DJX 0KQ

3UDK 095

2VIJ C44

3UDR 09F

3K5G BJC

3L3A 625

3RSV 3RS

3SKF PB7

2WF0 ZY0

4ACU QN7

3UDQ 09E

3KMY D8Y

3UDM 09A

3L38 879

3R2F PB0

3OOZ ZOO

2XFK AA9

3KN0 3TO

3K5F AYH

3K5D XLI

3IXK 929

4DPF 0LG

3OHF 3HF

3LPK Z76

3OHH 3HH

@<end>

mapk14

@<reference>

1ZYJ pinc_reference_files/mapk14/1ZYJ.mmtf BI5

@<align_prot>

1W83 L11

1BL7 SB4

1WBT WBT

2BAL PQA

@<align_sm_ligands>

3K3I JZJ

2YIS YIS

3KQ7 KQ7

4AA0 AA0

3LFC Z86

3L8X N4D

3HA8 5JZ

3FC1 52P

3FSF FSS

3FLN 3FN

4AAC AAV

3CTQ 337

3D83 GK6

3ZYA 2A8

3FLQ 891

3HL7 I47

3FMN 530

3FMK FMK

3S3I CQ0

3HRB I39

4A9Y RNY

4EWQ MWL

3GCS BAX

3HEC STI

3HP2 P36

3HEG BAX

3HVC GG5

3MPT 1GK

3FMJ FMJ

3OCG OCG

3FKL FKL

4AA4 QC0

2ZB1 GK4

3MVL 38P

2ZB0 GK3

3HV3 R49

3GI3 B10

3ZSI 52P

3IW6 PP0

3FL4 FL4

3NWW 3NW

3E92 G6A

3GFE P37

3HP5 52P

3GC7 B45

4AA5 NQB

3HUC G97

3BV2 P38

3FLS FLS

3FKN FKN

3D7Z GK5

3HLL I45

3FKO FKO

3FI4 FI4

3BX5 304

3FSK RO6

3FMM XI2

3IPH G11

3FLY FLY

3IW7 IPK

3FLZ FLZ

3E93 19B

3GCQ 1BU

3DT1 P40

3L8S BFF

3IW5 DF3

3FMH 533

3KF7 L9G

3DS6 A17

3FML FML

3LHJ LHJ

3BV3 P39

3FLW FLW

3ITZ P66

3HV4 L51

3ROC 29A

3HV6 R39

3MVM 39P

3HUB 469

3ZSG T75

3ZSH 469

3QUE 3FF

3S4Q NK0

2YIW YIW

3LFE Z84

3QUD N3F

3MW1 MIH

3ZS5 SB2

3PG3 DG7

3RIN I2O

2YIX YIX

3GCU R48

@<end>

cdk2

@<reference>

1H1P pinc_reference_files/cdk2/1H1P.mmtf CMG

@<align_prot>

1H0V UN4

1CKP PVB

1H1S 4SP

1JVP LIG

@<align_sm_ligands>

1PYE PM1

2VTJ LZ4

2UZL C94

1WCC CIG

2VTM LZM

3UNJ 0BX

2BHE BRY

1YKR 628

3PJ8 404

1PXI CK1

3FZ1 B98

2BPM 529

2C6M DT5

1VYW 292

3TNW F18

2UUE MTZ

2UZD C85

2DUV 371

2UZB C75

2VTN LZ7

2XNB Y8L

3EID PO5

1PXM CK5

2VTT LZD

2C69 CT8

3BHV VAR

2B53 D23

2VTI LZ3

2C5N CK8

1W0X OLO

3PXY JWS

2FVD LIA

3PXQ 2AN

2BTR U73

2R3Q 5SC

3LE6 2BZ

1URW I1P

2VU3 LZE

2R3M SCX

1PXJ CK2

3LFS A07

1Y8Y CT7

2UZN C96

2C6I DT1

2C5V CK4

2B55 D31

2VTA LZ1

3SW4 18K

2VTH LZ2

2W05 FRT

2VTR LZB

1PXN CK6

4ACM 7YG

1PXL CK4

2VTL LZ5

3MY5 RFZ

2W1H L0F

1VYZ N5B

2C6L DT4

2IW8 4SP

2R3O 2SC

2WIH P48

2C6K DT2

2R64 740

2R3F SC8

3LFN A27

1PXP CK8

2R3K SCQ

3EZV EZV

1PXO CK7

2R3N SCZ

3SW7 19K

1PXK CK3

1OIU N76

3NS9 NS9

2R3H SCE

2G9X NU5

2B52 D42

2WEV CK7

2C5Y MTW

2R3P 3SC

2VTQ LZA

2BKZ SBC

2BTS U32

3BHT MFR

3IG7 EFP

3EJ1 5BP

2C68 CT6

3UNK 0BY

3PXZ JWS

2C5X MTW

2BHH RYU

3IGG EFQ

1R78 FMD

2VTS LZC

2C6O 4SP

2C5O CK2

2UZE C95

2UZO C62

2W17 I19

2XMY CDK

2B54 D05

3LFQ A28

3BHU MHR

2EXM ZIP

2C6T DT5

2R3R 6SC

3EZR EZR

2A4L RRC

2R3G SC9

1W8C N69

3PY0 SU9

2VTO LZ8

2WIP P49

2C4G 514

2WPA 889

2R3J SCJ

3EOC T2A

2VTP LZ9

2J9M PY8

3S2P PMU

1Y91 CT9

2A0C CK9

2W06 FRV

3PY1 SU9

2CCH ATP

2I40 BLZ

@<end>

thrombin

@<reference>

1K21 pinc_reference_files/thrombin/1K21.mmtf IGN

@<align_prot>

1CA8 0KV

1DWB BEN

1BMN BM9

1D3P BT3

@<align_sm_ligands>

2CN0 F25

3RLY S29

2C8Z C2A

3RMO S04

1WBG L03

3BIV 11U

1OYT FSN

3C1K T15

2ZDA 32U

2C90 C1M

2ANK N12

1NZQ 162

2ZFP 19U

1YPJ UIB

1YPK CCR

3QWC 98P

1SB1 165

2ZG0 50U

1WAY L02

2ZC9 22U

1YPG UIR

3C27 DKK

1T4V 14A

2ZO3 33U

2CF9 348

3SHA P97

1YPM RA4

3SV2 P05

2BVR 4CP

2BXU C1D

1KTS C24

3RMN M41

1YPL RA8

2C93 C4M

1TA2 176

3TU7 0BM

1MU6 CDA

1VZQ SHY

2ZHQ 27U

1ZRB 062

3P17 99P

2ZDV 37U

2JH0 701

3PO1 MKY

2PKS G44

3T5F M34

1MU8 CDB

1O2G 696

3SI3 B03

2FEQ 34P

3QTO 10P

1NM6 L86

1O5G CR9

2ZHE 13U

3P70 BEN

2C8X C5M

2ZNK 31U

1SL3 170

1O0D 163

2BVS 2CE

2R2M I50

2V3H I25

3F68 91U

1NT1 T76

3DUX 64U

2BXT C2D

3QTV 06P

2BVX 5CB

2BDY UNB

2ZF0 51U

1KTT C02

1MUE CDD

1ZGI 382

2ANM CDO

2ZFR 46U

4AX9 N5N

1RIW OSC

3DT0 16U

2FES 3SP

3SHC B01

2A2X NA9

1TA6 177

3RMM M32

3BIU 10U

3PMH 0G7

2ZI2 24U

3DA9 44U

1YPE UIP

1T4U 81A

2JH5 895

2C8Y C3M

3EQ0 2TS

1ZGV 501

2CF8 ESH

3RML M31

2ZGX 29U

1XM1 GAH

2GDE SN3

2V3O I26

3RM2 S00

3RLW S28

1W7G MIU

2UUJ 896

2JH6 894

2ZGB 21U

2ZFQ 45U

2ZHF 49U

3QX5 02P

3LDX NLI

2ZHW 12U

3SI4 B04

1Z71 L17

2C8W C7M

2ZIQ 26U

@<end>

carbonicanhydrase2

@<reference>

1IF4 pinc_reference_files/carbonicanhydrase2/1IF4.mmtf FBS

@<align_prot>

1G52 F2B

1Z9Y FUN

1G1D FSB

1BN4 AL9

@<align_sm_ligands>

2HD6 BOS

3RYJ RYJ

3MHI J90

3S78 EVJ

2WD2 MS5

3OYQ OYQ

2POV I7B

3HKU TOR

3T5Z B09

2NNG ZYX

2POW I7C

3C7P POF

3HKQ 1SD

2WD3 MS4

3CAJ EZL

3DAZ MZM

3RYZ RYZ

3VBD 0FZ

2GD8 PO1

2WEO FBW

3MHM J75

3M5E JDR

3S8X E59

3BL0 BL0

3HKT 2SD

3SBH E65

3HFP MIZ

3BET CTF

3M04 BE9

3RZ7 RZ7

1ZH9 MPX

2EU3 FF3

3QYK IE2

2HNC 1SA

2FOS B17

2NNS M25

2Q1B LSA

2H15 B19

2FOU B22

2NNO M28

2EU2 5DS

2QOA MAJ

3FFP LC1

2POU I7A

2QP6 MB1

2HL4 BO1

3DD0 EZL

2AW1 COX

3HLJ V21

2QO8 3CC

2WEH FB1

2WEG FBV

2Q38 LSA

2WEJ FB2

3HS4 AZM

3M3X JS7

2HOC 1CN

2X7S WZC

3D9Z D9Z

3IGP DT7

2X7T WZB

2FOQ B15

3MHC ARZ

3M40 J45

3IBL O59

3MYQ E27

3MHL J71

3B4F TUO

2NNV M29

3K34 SUA

3MNU BON

3M2N J74

3M98 E02

2X7U WZA

3M1K BEW

3PO6 DT9

3N2P AYX

3M14 BEV

3IBU O48

3RYY RYY

3MNA DWH

3L14 I7B

3R16 5UN

3P5A IT2

3N4B WWZ

3N0N P9B

3R17 5UM

3M67 E36

3OY0 OY0

3MMF D9H

3RYV RYV

3MZC S6I

3M96 E38

3S75 EVG

3OKU VZ4

3MHO J43

3P5L IT5

3S77 EVI

3S73 EVF

3S76 EVH

3D8W D8W

3OIM VZ5

3KIG DA4

3S72 EVE

3RYX RYX

3S71 EVD

3N3J WWV

3RZ5 RZ5

3V5G 0F3

3T82 SG4

3S9T E49

3T5U A09

3SBI E90

3T85 SG7

3NB5 R21

3ML2 SU0

3SAX E50

3F4X KLT

3IBN O60

3T84 SG6

3IBI BOW

3OYS OYS

3RZ0 RZ0

3RZ1 RZ1

3V7X D7A

3T83 MG5

3RZ8 RZ8

3T83 SG5

@<end>

hivrt

@<reference>

1RT4 pinc_reference_files/hivrt/1RT4.mmtf UC1

@<align_prot>

1RTI HEF

1REV TB9

1RT1 MKC

1VRU AAP

@<align_sm_ligands>

3LAK KR1

3C6U M22

1TKZ H16

2HNZ PC0

1LWC NVP

1C1C 612

1LW0 NVP

2OPP HBQ

1TKT H12

1TL1 H18

1EP4 S11

1FK9 EFZ

1LWF NVP

1LW2 U05

1DTQ FPT

1LWE NVP

2RF2 MRX

1C1B GCA

1JLG UC1

2HNY NVP

1TKX GWB

1JLQ SBN

3DLE GFA

1DTT FTC

3DI6 PDZ

3E01 PZ2

3C6T M14

3FFI 3OB

2RKI TT1

3MEE T27

3I0R RT3

3LAL KRV

3DRP R8E

3DLG GWE

3MEC 65B

1TL3 H20

3M8Q DJZ

3QIP NVP

3LAN KBT

3T19 5MA

3M8P 65B

3LAM KRP

3NBP JGZ

3DYA PZL

3I0S RT7

2WON ZZE

@<end>

hivproteaseb1

@<reference>

1HVI pinc_reference_files/hivproteaseb1/1HVI.mmtf A77

@<align_prot>

1MET DMP

1HIH C20

1GNO U0E

1HVK A79

@<align_sm_ligands>

1A9M U0E

1YT9 OIS

1NPW LGZ

1G35 AHF

2FDE 385

1EC1 BEE

1EBY BEB

1W5X BE5

1T7K BH0

1AJX AH1

1AJV NMB

1W5W BE4

1NPA 3NH

1NPV L27

1D4H BEH

1EBW BEI

2AQU DR7

1ZSF 0ZS

2HS1 017

2I4U DJR

2BBB HH1

2IDW 017

1D4I BEG

1W5Y BE6

1DW6 0Q4

3NUO 478

1W5V BE3

1D4J MSC

1EC2 BEJ

1ZSR 0ZT

2I4X KGQ

2I4W KGQ

2I4V DJR

3GGV GGV

2I4D QFI

1EC0 BED

2UXZ HI1

3TL9 ROC

2UY0 HV1

3GGX GGX

3GGA GGW

@<end>

**Listing 5: Input for HAP 2 dataset**

@<reference>

2d59 hap2_reference_files/2d59.mmtf.gz

@<align_sm_ligands>

2d5a COA

@<end>

@<reference>

2iyt hap2_reference_files/2iyt.mmtf.gz

@<align_sm_ligands>

2iys SKM

@<end>

@<reference>

1alb hap2_reference_files/1alb.mmtf.gz

@<align_sm_ligands>

1lif STE

@<end>

@<reference>

1gqv hap2_reference_files/1gqv.mmtf.gz

@<align_sm_ligands>

2c05 B4P

@<end>

@<reference>

1eut hap2_reference_files/1eut.mmtf.gz

@<align_sm_ligands>

1eus DAN

@<end>

@<reference>

1uaj hap2_reference_files/1uaj.mmtf.gz

@<align_sm_ligands>

1uak SAM

@<end>

@<reference>

3c2e hap2_reference_files/3c2e.mmtf.gz

@<align_sm_ligands>

3c2o NTM

@<end>

@<reference>

1wu4 hap2_reference_files/1wu4.mmtf.gz

@<align_sm_ligands>

1wu5 XYP

@<end>

@<reference>

1quv hap2_reference_files/1quv.mmtf.gz

@<align_sm_ligands>

2brl POO

@<end>

@<reference>

1ia8 hap2_reference_files/1ia8.mmtf.gz

@<align_sm_ligands>

2r0u M54

@<end>

@<reference>

2qev hap2_reference_files/2qev.mmtf.gz

@<align_sm_ligands>

2pql TSS

@<end>

@<reference>

1ile hap2_reference_files/1ile.mmtf.gz

@<align_sm_ligands>

1jzs MRC

@<end>

@<reference>

2x16 hap2_reference_files/2x16.mmtf.gz

@<align_sm_ligands>

2x1t RES

@<end>

@<reference>

1fmk hap2_reference_files/1fmk.mmtf.gz

@<align_sm_ligands>

2h8h H8H

@<end>

@<reference>

1h7e hap2_reference_files/1h7e.mmtf.gz

@<align_sm_ligands>

1h7f C5P

@<end>

@<reference>

2exo hap2_reference_files/2exo.mmtf.gz

@<align_sm_ligands>

3cuf 9MR

@<end>

@<reference>

1a3h hap2_reference_files/1a3h.mmtf.gz

@<align_sm_ligands>

4a3h DCB

@<end>

@<reference>

2npo hap2_reference_files/2npo.mmtf.gz

@<align_sm_ligands>

3bfp FLC

@<end>

@<reference>

1kwb hap2_reference_files/1kwb.mmtf.gz

@<align_sm_ligands>

1kwc BPY

@<end>

@<reference>

2ohg hap2_reference_files/2ohg.mmtf.gz

@<align_sm_ligands>

2ohv NHL

@<end>

@<reference>

1f46 hap2_reference_files/1f46.mmtf.gz

@<align_sm_ligands>

1s1s WAC

@<end>

@<reference>

2jdw hap2_reference_files/2jdw.mmtf.gz

@<align_sm_ligands>

7jdw DAV

@<end>

@<reference>

1pud hap2_reference_files/1pud.mmtf.gz

@<align_sm_ligands>

1k4h APQ

@<end>

@<reference>

1izd hap2_reference_files/1izd.mmtf.gz

@<align_sm_ligands>

1ize MAN

@<end>

@<reference>

3acb hap2_reference_files/3acb.mmtf.gz

@<align_sm_ligands>

3acd IMP

@<end>

@<reference>

1bsq hap2_reference_files/1bsq.mmtf.gz

@<align_sm_ligands>

1gx9 REA

@<end>

@<reference>

1brq hap2_reference_files/1brq.mmtf.gz

@<align_sm_ligands>

1rbp RTL

@<end>

@<reference>

1i76 hap2_reference_files/1i76.mmtf.gz

@<align_sm_ligands>

1bzs BSI

@<end>

@<reference>

2hc1 hap2_reference_files/2hc1.mmtf.gz

@<align_sm_ligands>

2h03 3UN

@<end>

@<reference>

1py3 hap2_reference_files/1py3.mmtf.gz

@<align_sm_ligands>

3d5i SGP

@<end>

@<reference>

2dul hap2_reference_files/2dul.mmtf.gz

@<align_sm_ligands>

2ejt SAM

@<end>

@<reference>

1lcl hap2_reference_files/1lcl.mmtf.gz

@<align_sm_ligands>

1qkq MAN

@<end>

@<reference>

2j1x hap2_reference_files/2j1x.mmtf.gz

@<align_sm_ligands>

2x0u X0U

@<end>

@<reference>

1rbb hap2_reference_files/1rbb.mmtf.gz

@<align_sm_ligands>

1eos U2G

@<end>

@<reference>

1t80 hap2_reference_files/1t80.mmtf.gz

@<align_sm_ligands>

1t7v NAG

@<end>

@<reference>

1eyp hap2_reference_files/1eyp.mmtf.gz

@<align_sm_ligands>

1fm8 DDC

@<end>

@<reference>

1cz1 hap2_reference_files/1cz1.mmtf.gz

@<align_sm_ligands>

1eqc CTS

@<end>

@<reference>

2cc9 hap2_reference_files/2cc9.mmtf.gz

@<align_sm_ligands>

2ccb RBF

@<end>

@<reference>

3erf hap2_reference_files/3erf.mmtf.gz

@<align_sm_ligands>

3ibh GSH

@<end>

@<reference>

1b73 hap2_reference_files/1b73.mmtf.gz

@<align_sm_ligands>

1b74 DGN

@<end>

\# TODO choose here (only lignad is common peptide)

@<reference>

1tje hap2_reference_files/1tje.mmtf.gz

@<align_non_sm_ligands>

1tke SER-A-500

@<end>

@<reference>

6rhn hap2_reference_files/6rhn.mmtf.gz

@<align_sm_ligands>

5rhn 8BR

@<end>

@<reference>

1pjb hap2_reference_files/1pjb.mmtf.gz

@<align_sm_ligands>

1pjc NAD

@<end>

@<reference>

2qbv hap2_reference_files/2qbv.mmtf.gz

@<align_sm_ligands>

2vkl MLT

@<end>

@<reference>

1jam hap2_reference_files/1jam.mmtf.gz

@<align_sm_ligands>

2oxd K32

@<end>

@<reference>

1sjs hap2_reference_files/1sjs.mmtf.gz

@<align_sm_ligands>

9icd NAP

@<end>

@<reference>

1egu hap2_reference_files/1egu.mmtf.gz

@<align_sm_ligands>

1f9g ASC

@<end>

@<reference>

2r5g hap2_reference_files/2r5g.mmtf.gz

@<align_sm_ligands>

2per MNB

@<end>

\# TODO choose here (ligand is common peptide)

@<reference>

3htz hap2_reference_files/3htz.mmtf.gz

@<align_non_sm_ligands>

1ygb SER-A-501

@<end>

@<reference>

2wbz hap2_reference_files/2wbz.mmtf.gz

@<align_sm_ligands>

1lxz TLA

@<end>

@<reference>

1gy0 hap2_reference_files/1gy0.mmtf.gz

@<align_sm_ligands>

1og1 TAD

@<end>

@<reference>

1b8p hap2_reference_files/1b8p.mmtf.gz

@<align_sm_ligands>

1b8v NAD

@<end>

@<reference>

1qba hap2_reference_files/1qba.mmtf.gz

@<align_sm_ligands>

1qbb CBS

@<end>

@<reference>

2ebg hap2_reference_files/2ebg.mmtf.gz

@<align_sm_ligands>

2ei5 BTB

@<end>

@<reference>

1nie hap2_reference_files/1nie.mmtf.gz

@<align_sm_ligands>

2bw5 MLI

@<end>

@<reference>

1lz7 hap2_reference_files/1lz7.mmtf.gz

@<align_sm_ligands>

1ziz GAL

@<end>

@<reference>

1rtc hap2_reference_files/1rtc.mmtf.gz

@<align_sm_ligands>

1br5 NEO

@<end>

@<reference>

2zco hap2_reference_files/2zco.mmtf.gz

@<align_sm_ligands>

2zy1 830

@<end>

@<reference>

1gbs hap2_reference_files/1gbs.mmtf.gz

@<align_sm_ligands>

1lsp BUL

@<end>

@<reference>

3go7 hap2_reference_files/3go7.mmtf.gz

@<align_sm_ligands>

3go6 RIB

@<end>

@<reference>

2bls hap2_reference_files/2bls.mmtf.gz

@<align_sm_ligands>

3gr2 GF4

@<end>

@<reference>

1kxo hap2_reference_files/1kxo.mmtf.gz

@<align_sm_ligands>

1lnm DTX

@<end>

@<reference>

3gvr hap2_reference_files/3gvr.mmtf.gz

@<align_sm_ligands>

1r3y CP3

@<end>

@<reference>

1dhn hap2_reference_files/1dhn.mmtf.gz

@<align_sm_ligands>

1rsi 977

@<end>

@<reference>

1dvq hap2_reference_files/1dvq.mmtf.gz

@<align_sm_ligands>

1dvy BPD

@<end>

@<reference>

1x1h hap2_reference_files/1x1h.mmtf.gz

@<align_sm_ligands>

1x1j 46D

@<end>

@<reference>

2ris hap2_reference_files/2ris.mmtf.gz

@<align_sm_ligands>

2riu 5RP

@<end>

@<reference>

1hcl hap2_reference_files/1hcl.mmtf.gz

@<align_sm_ligands>

1aq1 STU

@<end>

@<reference>

2cl3 hap2_reference_files/2cl3.mmtf.gz

@<align_sm_ligands>

3bho B4P

@<end>

@<reference>

1v8i hap2_reference_files/1v8i.mmtf.gz

@<align_sm_ligands>

1v8l APR

@<end>

@<reference>

1nx2 hap2_reference_files/1nx2.mmtf.gz

@<align_sm_ligands>

1nx3 ISA

@<end>

@<reference>

1wka hap2_reference_files/1wka.mmtf.gz

@<align_sm_ligands>

1wk9 TSB

@<end>

@<reference>

1e8y hap2_reference_files/1e8y.mmtf.gz

@<align_sm_ligands>

1e8z STU

@<end>

@<reference>

1jsm hap2_reference_files/1jsm.mmtf.gz

@<align_sm_ligands>

1jso NAG

@<end>

@<reference>

1qid hap2_reference_files/1qid.mmtf.gz

@<align_sm_ligands>

1qti GNT

@<end>

@<reference>

1tpo hap2_reference_files/1tpo.mmtf.gz

@<align_sm_ligands>

1o2i 655

@<end>

@<reference>

6yas hap2_reference_files/6yas.mmtf.gz

@<align_sm_ligands>

5yas FAC

@<end>

@<reference>

1pjb hap2_reference_files/1pjb.mmtf.gz

@<align_sm_ligands>

1say PYR

@<end>

\# TODO choose here (NAG removed)

@<reference>

1bsi hap2_reference_files/1bsi.mmtf.gz

@<align_sm_ligands>

1xcw 3SA

@<end>

@<reference>

2psr hap2_reference_files/2psr.mmtf.gz

@<align_sm_ligands>

2wor 2AN

@<end>

@<reference>

3aap hap2_reference_files/3aap.mmtf.gz

@<align_sm_ligands>

3aar ANP

@<end>

@<reference>

16gs hap2_reference_files/16gs.mmtf.gz

@<align_sm_ligands>

3csj CBL

@<end>

@<reference>

1ks9 hap2_reference_files/1ks9.mmtf.gz

@<align_sm_ligands>

1yon A2R

@<end>

@<reference>

3f9q hap2_reference_files/3f9q.mmtf.gz

@<align_sm_ligands>

1lee R36

@<end>

@<reference>

2ozq hap2_reference_files/2ozq.mmtf.gz

@<align_sm_ligands>

1znl DE1

@<end>

@<reference>

2wxr hap2_reference_files/2wxr.mmtf.gz

@<align_sm_ligands>

2wxo ZZP

@<end>

@<reference>

1eut hap2_reference_files/1eut.mmtf.gz

@<align_sm_ligands>

1euu GAL

@<end>

@<reference>

1jks hap2_reference_files/1jks.mmtf.gz

@<align_sm_ligands>

3eha ANP

@<end>

@<reference>

3gez hap2_reference_files/3gez.mmtf.gz

@<align_sm_ligands>

3gf2 SAL

@<end>

@<reference>

1ufk hap2_reference_files/1ufk.mmtf.gz

@<align_sm_ligands>

2zbr SFG

@<end>

@<reference>

2hjw hap2_reference_files/2hjw.mmtf.gz

@<align_sm_ligands>

3jrx S1A

@<end>

@<reference>

2w91 hap2_reference_files/2w91.mmtf.gz

@<align_sm_ligands>

2w92 NGT

@<end>

@<reference>

5dfr hap2_reference_files/5dfr.mmtf.gz

@<align_sm_ligands>

1ra1 NAP

@<end>

@<reference>

1nux hap2_reference_files/1nux.mmtf.gz

@<align_sm_ligands>

1cnq F6P

@<end>

@<reference>

1doz hap2_reference_files/1doz.mmtf.gz

@<align_sm_ligands>

1c1h MMP

@<end>

@<reference>

3ba1 hap2_reference_files/3ba1.mmtf.gz

@<align_sm_ligands>

3baz NAP

@<end>

@<reference>

1tqi hap2_reference_files/1tqi.mmtf.gz

@<align_sm_ligands>

1tqm ANP

@<end>

\# TODO choose here (removed LLP)

@<reference>

2gpn hap2_reference_files/2gpn.mmtf.gz

@<align_sm_ligands>

1z6p 194

@<end>

\# TODO choose here (lignad is common peptide)

@<reference>

1iiw hap2_reference_files/1iiw.mmtf.gz

@<align_non_sm_ligands>

1ii5 GLU-A-999

@<end>

@<reference>

1bk7 hap2_reference_files/1bk7.mmtf.gz

@<align_sm_ligands>

1uca U2P

@<end>

@<reference>

1s4q hap2_reference_files/1s4q.mmtf.gz

@<align_sm_ligands>

1znz GDP

@<end>

@<reference>

4pgm hap2_reference_files/4pgm.mmtf.gz

@<align_sm_ligands>

1bq4 BHC

@<end>

@<reference>

1ri5 hap2_reference_files/1ri5.mmtf.gz

@<align_sm_ligands>

1ri4 SAM

@<end>

@<reference>

2cpl hap2_reference_files/2cpl.mmtf.gz

@<align_sm_ligands>

1w8l 1P3

@<end>

@<reference>

1sca hap2_reference_files/1sca.mmtf.gz

@<align_sm_ligands>

1bfu DIO

@<end>

@<reference>

1xqz hap2_reference_files/1xqz.mmtf.gz

@<align_sm_ligands>

1yxt ANP

@<end>

@<reference>

1gta hap2_reference_files/1gta.mmtf.gz

@<align_sm_ligands>

1m99 GTS

@<end>

\# TODO choose here (removed IMD)

@<reference>

1jrl hap2_reference_files/1jrl.mmtf.gz

@<align_sm_ligands>

1v2g OCA

@<end>

@<reference>

1qtr hap2_reference_files/1qtr.mmtf.gz

@<align_sm_ligands>

1x2e ATX

@<end>

@<reference>

2wxr hap2_reference_files/2wxr.mmtf.gz

@<align_sm_ligands>

2wxh ZZO

@<end>

@<reference>

2cwk hap2_reference_files/2cwk.mmtf.gz

@<align_sm_ligands>

2dxf GNP

@<end>

@<reference>

1erk hap2_reference_files/1erk.mmtf.gz

@<align_sm_ligands>

3erk SB4

@<end>

@<reference>

3dw3 hap2_reference_files/3dw3.mmtf.gz

@<align_sm_ligands>

1oyo 3ID

@<end>

@<reference>

2bv9 hap2_reference_files/2bv9.mmtf.gz

@<align_sm_ligands>

2bvd ISX

@<end>

\# TODO choose here (removed NAG)

@<reference>

1gpi hap2_reference_files/1gpi.mmtf.gz

@<align_sm_ligands>

1z3w IDC

@<end>

@<reference>

2qht hap2_reference_files/2qht.mmtf.gz

@<align_sm_ligands>

2qhs OCA

@<end>

@<reference>

3pte hap2_reference_files/3pte.mmtf.gz

@<align_sm_ligands>

1ikg REX

@<end>

\# TODO choose here (removed NAG)

@<reference>

1j1q hap2_reference_files/1j1q.mmtf.gz

@<align_sm_ligands>

1j1s FMP

@<end>

@<reference>

3fw6 hap2_reference_files/3fw6.mmtf.gz

@<align_sm_ligands>

3ii1 BGC

@<end>

@<reference>

1dup hap2_reference_files/1dup.mmtf.gz

@<align_sm_ligands>

1dud DUD

@<end>

@<reference>

1vp3 hap2_reference_files/1vp3.mmtf.gz

@<align_sm_ligands>

1jsz NDM

@<end>

@<reference>

2vef hap2_reference_files/2vef.mmtf.gz

@<align_sm_ligands>

2veg PMM

@<end>

@<reference>

3iep hap2_reference_files/3iep.mmtf.gz

@<align_sm_ligands>

3ier PG4

@<end>

@<reference>

2zgl hap2_reference_files/2zgl.mmtf.gz

@<align_sm_ligands>

2zgn GAL

@<end>

@<reference>

1tw7 hap2_reference_files/1tw7.mmtf.gz

@<align_sm_ligands>

1rpi GLC

@<end>

@<reference>

3ewq hap2_reference_files/3ewq.mmtf.gz

@<align_sm_ligands>

3ewr APR

@<end>

@<reference>

1gcg hap2_reference_files/1gcg.mmtf.gz

@<align_sm_ligands>

1gca GAL

@<end>

@<reference>

3cj1 hap2_reference_files/3cj1.mmtf.gz

@<align_sm_ligands>

3cj7 AMP

@<end>

@<reference>

1pah hap2_reference_files/1pah.mmtf.gz

@<align_sm_ligands>

1lrm HBI

@<end>

@<reference>

1nm8 hap2_reference_files/1nm8.mmtf.gz

@<align_sm_ligands>

1s5o 152

@<end>

@<reference>

2d2r hap2_reference_files/2d2r.mmtf.gz

@<align_sm_ligands>

2dtn DPO

@<end>

@<reference>

3h97 hap2_reference_files/3h97.mmtf.gz

@<align_sm_ligands>

3h9b NOT

@<end>

@<reference>

3a7f hap2_reference_files/3a7f.mmtf.gz

@<align_sm_ligands>

3a7i ADE

@<end>

\# TODO choose here (removed TLA)

@<reference>

1frz hap2_reference_files/1frz.mmtf.gz

@<align_sm_ligands>

1fs5 16G

@<end>

@<reference>

2wck hap2_reference_files/2wck.mmtf.gz

@<align_sm_ligands>

2wcj M21

@<end>

@<reference>

1gta hap2_reference_files/1gta.mmtf.gz

@<align_sm_ligands>

1m9a GTX

@<end>

@<reference>

2bgt hap2_reference_files/2bgt.mmtf.gz

@<align_sm_ligands>

1qkj UDP

@<end>

@<reference>

2g52 hap2_reference_files/2g52.mmtf.gz

@<align_sm_ligands>

1pq5 ARG

@<end>

@<reference>

2fya hap2_reference_files/2fya.mmtf.gz

@<align_sm_ligands>

2fy7 PGE

@<end>

@<reference>

2gfv hap2_reference_files/2gfv.mmtf.gz

@<align_sm_ligands>

3g11 P9C

@<end>

@<reference>

1jud hap2_reference_files/1jud.mmtf.gz

@<align_sm_ligands>

1qh9 LAC

@<end>

\# TODO choose here (lignad is common peptide)

@<reference>

1y2q hap2_reference_files/1y2q.mmtf.gz

@<align_non_sm_ligands>

2hkz SER-A-201

@<end>

@<reference>

1s7k hap2_reference_files/1s7k.mmtf.gz

@<align_sm_ligands>

1s7f MLA

@<end>

@<reference>

1j4b hap2_reference_files/1j4b.mmtf.gz

@<align_sm_ligands>

1loo GTP

@<end>

@<reference>

3ado hap2_reference_files/3ado.mmtf.gz

@<align_sm_ligands>

3adp NAI

@<end>

\# TODO choose here (removed GOL)

@<reference>

2vk5 hap2_reference_files/2vk5.mmtf.gz

@<align_sm_ligands>

2bf6 SIA

@<end>

@<reference>

1sgk hap2_reference_files/1sgk.mmtf.gz

@<align_sm_ligands>

1dtp APU

@<end>

@<reference>

226l hap2_reference_files/226l.mmtf.gz

@<align_sm_ligands>

225l PXY

@<end>

\# TODO choose here (removed HAI)

@<reference>

2h4x hap2_reference_files/2h4x.mmtf.gz

@<align_sm_ligands>

2h52 3PG

@<end>

@<reference>

3jyl hap2_reference_files/3jyl.mmtf.gz

@<align_sm_ligands>

3jyn NDP

@<end>

@<reference>

3wrp hap2_reference_files/3wrp.mmtf.gz

@<align_sm_ligands>

1wrp TRP

@<end>

@<reference>

2fgz hap2_reference_files/2fgz.mmtf.gz

@<align_sm_ligands>

2fh6 GLC

@<end>

@<reference>

1lp8 hap2_reference_files/1lp8.mmtf.gz

@<align_sm_ligands>

1lpc CMP

@<end>

@<reference>

1cy2 hap2_reference_files/1cy2.mmtf.gz

@<align_sm_ligands>

1cy7 TMP

@<end>

@<reference>

1ahc hap2_reference_files/1ahc.mmtf.gz

@<align_sm_ligands>

1aha ADE

@<end>

\# TODO choose here (removed GDP)

@<reference>

2a78 hap2_reference_files/2a78.mmtf.gz

@<align_sm_ligands>

2a9k NAD

@<end>

@<reference>

2ys7 hap2_reference_files/2ys7.mmtf.gz

@<align_sm_ligands>

2yrx AMP

@<end>

@<reference>

1zn9 hap2_reference_files/1zn9.mmtf.gz

@<align_sm_ligands>

1zn8 AMP

@<end>

@<reference>

1akz hap2_reference_files/1akz.mmtf.gz

@<align_sm_ligands>

3fci 3FI

@<end>

@<reference>

2o9p hap2_reference_files/2o9p.mmtf.gz

@<align_sm_ligands>

2z1s CTT

@<end>

@<reference>

1uyl hap2_reference_files/1uyl.mmtf.gz

@<align_sm_ligands>

2byi 2DD

@<end>

\# TODO choose here (removed IMD)

@<reference>

3ftv hap2_reference_files/3ftv.mmtf.gz

@<align_sm_ligands>

3ftw 11X

@<end>

@<reference>

1zhf hap2_reference_files/1zhf.mmtf.gz

@<align_sm_ligands>

1zg3 2HI

@<end>

@<reference>

2uyo hap2_reference_files/2uyo.mmtf.gz

@<align_sm_ligands>

2uyq SAM

@<end>

@<reference>

2paw hap2_reference_files/2paw.mmtf.gz

@<align_sm_ligands>

1pax DHQ

@<end>

@<reference>

1wos hap2_reference_files/1wos.mmtf.gz

@<align_sm_ligands>

1wor RED

@<end>

\# TODO choose here (removed BMA, MAN, NAG)

@<reference>

2qwa hap2_reference_files/2qwa.mmtf.gz

@<align_sm_ligands>

2qwf G20

@<end>

@<reference>

1adi hap2_reference_files/1adi.mmtf.gz

@<align_sm_ligands>

1hon GNH

@<end>

@<reference>

2r60 hap2_reference_files/2r60.mmtf.gz

@<align_sm_ligands>

2r68 SUP

@<end>

@<reference>

2btz hap2_reference_files/2btz.mmtf.gz

@<align_sm_ligands>

2bu7 TF3

@<end>

@<reference>

2j64 hap2_reference_files/2j64.mmtf.gz

@<align_sm_ligands>

2j5z GAL

@<end>

@<reference>

1iq8 hap2_reference_files/1iq8.mmtf.gz

@<align_sm_ligands>

1it8 PQ0

@<end>

\# TODO choose here (removed GAL, NGA)

@<reference>

1y2v hap2_reference_files/1y2v.mmtf.gz

@<align_sm_ligands>

1y2w NAG

@<end>

\# TODO choose here (removed MPD)

@<reference>

1kwg hap2_reference_files/1kwg.mmtf.gz

@<align_sm_ligands>

1kwk GAL

@<end>

@<reference>

1s0d hap2_reference_files/1s0d.mmtf.gz

@<align_sm_ligands>

1i1e DM2

@<end>

@<reference>

1ctq hap2_reference_files/1ctq.mmtf.gz

@<align_sm_ligands>

1p2s GNP

@<end>

@<reference>

3f6f hap2_reference_files/3f6f.mmtf.gz

@<align_sm_ligands>

3gh6 GSH

@<end>

@<reference>

1khg hap2_reference_files/1khg.mmtf.gz

@<align_sm_ligands>

1m51 TSX

@<end>

@<reference>

1sll hap2_reference_files/1sll.mmtf.gz

@<align_sm_ligands>

3sli SKD

@<end>

@<reference>

1x6l hap2_reference_files/1x6l.mmtf.gz

@<align_sm_ligands>

1x6n AO3

@<end>

**Listing 6: Input for DUD-E dataset**

@<no_align_sm_ligands>

3eml ZMA

2hzi JIN

3bkl KAW

1e66 HUX

2e1w FR6

2oi0 283

2vt4 P32

3ny8 JRZ

3cqw CQW

3d0e G93

2hv5 ZST

1l2s STC

2am9 TES

1s3b RMA

3l5d BDV

3d4q SM5

1bcd FMS

2cnk MY2

1h00 FAP

3bwm DNC

1r9o FLP

3nxu RIT

3krj KRJ

3odu ITD

1lru BB2

3frj A49

2i78 KIQ

3pbl ETQ

3nxo D2B

2rgp HYZ

1sj0 E4D

2fsz OHT

3kl6 443

1w7x 413

2nnq T4B

3bz3 YAM

3c4f C4F

1j4h SUB

3e37 ED5

1zw5 ZOL

3bqd DAY

2v3f BTB

3kgc ZK1

1vso AT1

3max LLX

3f07 AGE

3nf7 CIW

1xl2 189

3lan KBT

3ccw 4HI

1uyg PU2

3f9m MRK

2oj9 BMI

2h7l 665

2ica 2IC

3lpb NVB

3cjo K30

3g0e B49

2b8t THM

2i0e PDS

2of2 547

3chp 4BO

3m2w L8I

2aa2 AS4

3lq8 88Z

2ojg 19A

2zdt 46C

2qd9 LGF

830c RS1

3eqh 5BM

1qw6 3AR

1b9v RA2

1kvo OAP

3l3m A92

1udt VIA

2oyu IMS

3ln1 CEL

2owb 626

3bgs DIH

2p54 735

2znp K55

2gtk 208

3kba WOW

2azr 982

1njs KEU

1c8k CPB

1d3g BRE

3g6z A7T

2etr Y27

1mv9 HXA

1li4 NOC

3el8 PD5

3hmm 855

1qx4 G24

1ype UIP

2ayw ONO

2zec 11N

1syn F89

1sqt UI3

2p2i 608

3biz 61E

3hl5 9JZ

**Listing 7: Input for CSAR dataset**

@<no_align_sm_ligands>

4FW3 L52

4fW4 3P3

4FW5 L58

4FW6 L59

4FW7 L63

4EK4 1CK

4EK5 03K

4FKI 09K

4EK6 10K

4FJK DTP

4FKL C2K

4EK8 16K

3SW4 18K

3SW7 19K

4FKO 20K

4FKP LS5

4FKQ 42K

4FKR 45K

4FKS 46K

4FKT 48K

4FKU 60K

4FKV 61K

4FKW 62K

4FX3 60K
